# Age-specific genomic and transcriptomic variation reveals limited evidence for *cis-*regulatory interactions modulating aging in *Saccharomyces cerevisiae*

**DOI:** 10.64898/2025.12.12.689579

**Authors:** Kaitlin M McHugh, Felipe S Barreto, Molly K Burke

**Author notes:** Authors for correspondence: Kaitlin M. McHugh and Molly K. Burke, Department of Integrative Biology, Oregon State University, Corvallis, OR, USA, or, fax: NA.

## Abstract

The budding yeast *Saccharomyces cerevisiae* is a well-established model for studying the genetic basis of complex traits, and it is a powerful system for investigating mechanisms of aging. Here, we examine the genomic and transcriptomic factors contributing to increased replicative age in recombinant yeast populations harboring standing genetic variation. Using Fluorescence-Activated Cell Sorting (FACS), we isolated young and aged cohort pairs across twelve biological replicates and sequenced their progeny to assess patterns of differentiation at the nucleotide and transcription levels. Most differentiated alleles were located in coding regions, including significant variants within 132 unique genes. Transcriptomic analysis revealed 60 differentially expressed genes in aged populations, including 18 genes with increased expression in aged cohorts, and 42 genes with decreased expression. Although only two genes (*RFA3* and *WSC4)* were implicated in both genomic and transcriptomic analyses, functional overlap associated with protein homeostasis, DNA repair, and cell cycle regulation was evident across datasets. Notably, we found no strong evidence that differentially expressed genes were more likely to occur in close proximity to significant gene variants. This suggests that late-life survival is not predominantly governed by local *cis*-regulatory interactions (e.g. variants within or near coding regions). These findings underscore the power of integrating genomic and transcriptomic data to elucidate the genetics of complex traits such as aging, demonstrating how multi-omics approaches can reveal functional relationships that may be overlooked by single-layer analyses.

**Significance Statement:** While many individual genes contributing to aging and lifespan have been identified, our understanding of the polygenic interactions and regulatory processes that contribute to phenotypic variation in these traits is much more limited. Using a recombinant population of yeast, we identify novel links between genetic variation and the phenotype of replicative age. Additionally, we find little evidence for local *cis*-regulatory interactions, suggesting that downstream regulation or *trans*-regulatory processes may serve more dominant roles in modulating aging. These results reveal new insights into the role of polygenicity in the evolution and regulation of age-associated phenotypes.

## Introduction

Aging is a nearly universal process, yet the mechanisms that regulate its onset and progression remain incompletely understood. Identifying the genes and pathways that determine the timing and severity of aging offers the possibility of predicting or even delaying its physiological consequences. While many evolutionarily conserved regulators of aging have been described (López-Otín et al. 2013; Lemoine 2021; López-Otín et al. 2023), less is known about how sequence-level variation alters gene expression to shape aging. Deeper insight into this process could shed light on how natural genetic variation contributes to the evolution of aging phenotypes.

Gene expression is governed by both *cis-* regulatory and *trans-*regulatory elements. *Cis-*regulatory elements (e.g., promoters, enhancers, silencers) typically control the activity of linked genes through direct physical interactions, whereas *trans-*regulatory elements (e.g., transcription factors, regulatory RNAs) regulate unlinked genes via diffusion-mediated interactions. *Cis*-regulatory changes are of particular evolutionary interest because they are thought to contribute disproportionately to adaptive divergence (Wittkopp & Kalay 2011). Accordingly, dissecting the relative contributions of *cis*- and *trans*-acting factors provides a powerful framework for evaluating how genetic variation shapes aging-related phenotypes, by way of gene expression.

In theory, integrating genome sequence data with gene expression profiles of individuals from phenotypically diverse populations offers a means to distinguish *cis*- from *trans*-acting regulatory effects (Li & Ritchie 2021). Given the scale necessary to achieve this goal of integrating data across phenotypic, genomic and transcriptomic levels, laboratory studies with model organisms are especially useful. Studies with this objective have successfully identified regulatory variants underlying complex traits such as thermotolerance (Lecheta et al. 2020) and metabolic responses (Cubillos et al. 2017; Franco-Duarte et al. 2017), but applications to aging remain comparatively rare.

The yeast *Saccharomyces cerevisiae* has long been recognized as a powerful model system for aging research. Yeast share many conserved pathways with higher eukaryotes, and hundreds of genes have interchangeable human orthologs (Kachroo et al. 2015; Skrzypek et al. 2018). Replicative lifespan, defined as the total number of asexual divisions a cell undergoes, serves as a widely used proxy for aging in dividing cells and parallels aspects of human cellular senescence (Janssens & Veenhoff 2016). The replicative age of yeast mother cells can be measured via bud-scar staining, and fluorescence-activated cell sorting (FACS) enables separation of young and old cohorts for molecular profiling (Chen & Contreras 2004; Hendrickson et al. 2018). These features, combined with extensive genomic resources, make *S. cerevisiae* an especially tractable system for dissecting the genetic regulation of aging.

Here we leverage an outbred yeast population (“4S”; Phillips et al. 2021) generated from repeated outcrossing of four diverse ancestral strains. This population maintains extensive heterozygosity and standing genetic variation, providing a unique opportunity to evaluate how *cis-* and *trans-*regulatory elements contribute to expression differences associated with replicative aging. By stratifying cells by age and combining pooled-population sequencing with transcriptomic profiling, we identify candidate regulatory variants that underlie differential expression in aged cohorts. Our analyses reveal both known and novel genetic contributors to longevity-associated pathways, including genes involved in genome stability, protein homeostasis, epigenetic modifications, and nutrient signaling. However, we find limited evidence for an overrepresentation of local *cis*-regulatory effects, suggesting that longevity in this population is shaped primarily by complex, non-local regulatory interactions.

## Results

### Allele and haplotype frequency differentiation

We identified 67,049 biallelic SNPs that met our filtering criteria. Genome-wide coverage across all replicates averaged 117x (range: 40x-277x; Supplementary Table 1). Additionally, we identified 5,129 insertions/deletions (indels) that met filtering criteria with an average genome-wide coverage of 116 (range: 36x-278x; Supplementary Table 1). After applying genome-wide thresholds, we identified a total of 669 SNPs and 132 indels with significant difference in frequency between conditions (Figure 1A, Supplementary Figure 1). The complete list of significant SNPs and indels were located within 233 unique genomic features. Out of the original 801 variants, 463 were located within 132 unique genes, which we refer to as our “candidate genomic variant list” (Supplementary Table 2). We utilized SNPeff to find the gene feature most closely associated with each SNP. Two of the indels were not annotated within the SNPeff tool, but were noted as being within an autonomously replicating sequence (*ARS319*) in the UCSC genome browser. The resulting list of gene features was used for GO-term enrichment analysis, which detected functions associated with glycoprotein metabolism and the cell periphery. (Figure 2, Supplementary Table 3).

**Figure 1:**
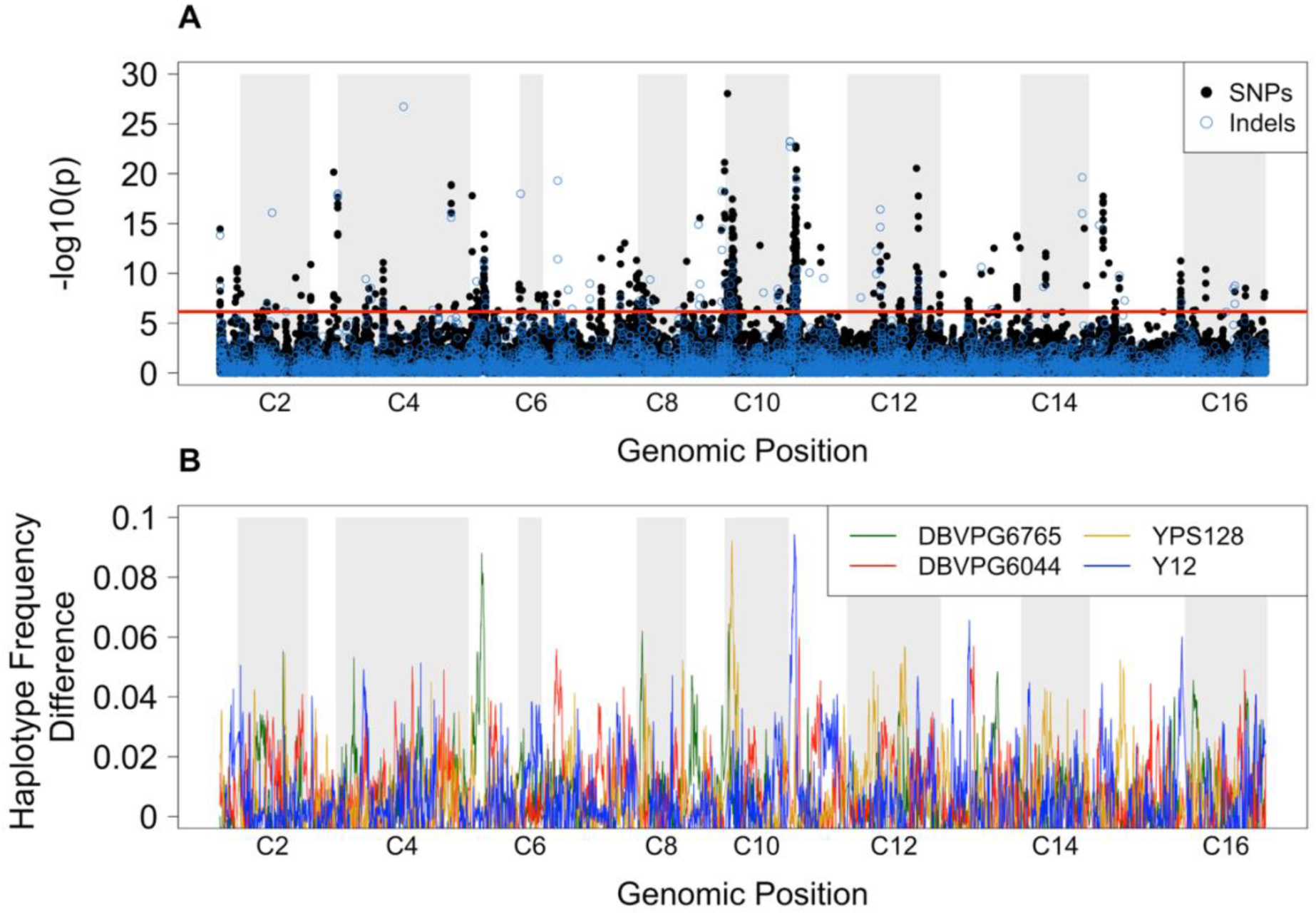
Gene variant frequency differentiation across the genome. **(A)** a paired CMH-test of SNP and indel frequencies across the genome reveals regions of significant differentiation. The red line indicates a Bonferroni corrected threshold of α=0.05 differentiating the most significant variants. **(B)** A sliding-window analysis of variation in ancestral haplotype frequencies between young and aged cohorts shows three primary regions of the genome in which a single ancestral haplotype is favored.

**Figure 2:**
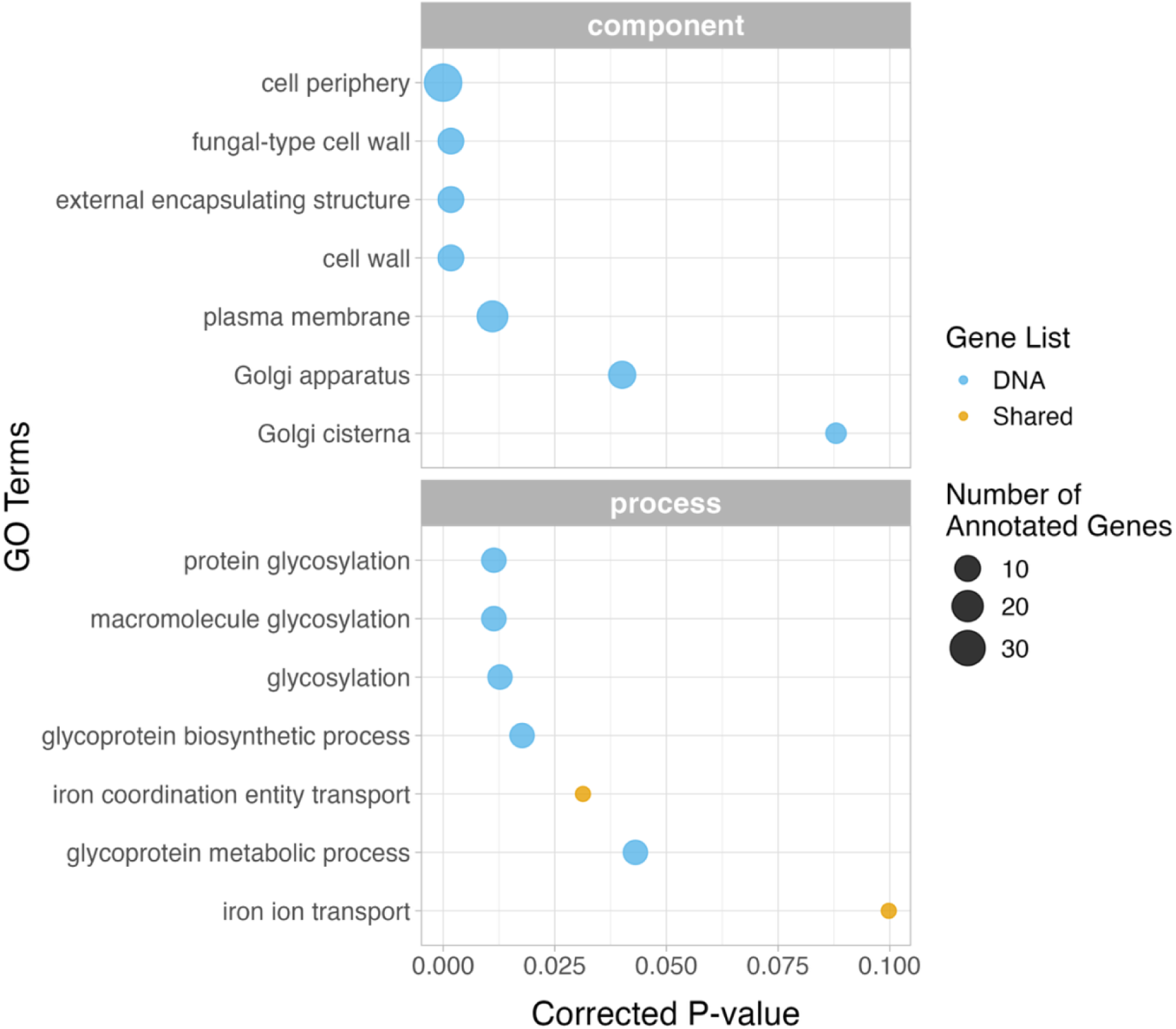
GO-term analysis for the genomic and transcriptomic gene lists. GO-term analyses were conducted separately for the genomic variant list, and the shared list of genes found in both genomic and transcriptomic variant lists. No significant terms were found for the transcriptomic variant list alone.

Haplotype-level analyses revealed distinct ancestral backgrounds associated with regions of the genome harboring significant SNPs (Figure 1B). Visual comparisons of Figures 1A and 1B led to the designation of three main peaks of interest, on chromosomes 5, 10, and 11. In addition to surpassing the threshold in Figure 1A, each of these peaks shows a single ancestral haplotype rising to prominence (Figure 1B) and the average percent divergence between all haplotypes is greater than 6% (Supplementary Figure 2). Notably, each of these peaks originated from a different ancestral haplotype; the DBVPG6765 European wine strain was elevated in the chromosome 5 peak, the YPS128 North American wild strain was isolated in the chromosome 10 peak, and the Y12 Japanese sake strain was elevated in the chromosome 11 peak (Supplementary Table 4). Additionally, several potentially interesting features were identified under each significant peak. Chromosome 5 included variants within *MNN1, YND1, FMP52, NOP16*. In chromosome 10, the most significant genic variants were found within *IMA5, MNN5, SWI3, BBC1,* and *RFA3*. The most significant genic variants within chromosome 11 included *PTK1, MNN4, TOR2*, and *YKL202W*. Peaks in each of these three chromosomes also contained a number of non-genic significant variants found within 5kb of an annotated gene.

### Differential gene expression between aged and young cohorts

Sequencing of RNA-seq libraries yielded an average of 21.8 million reads after quality trimming, and 16.5 million counted per sample after mapping (Supplementary Table 5). Mapping rates ranged from 57.3% to 80.1%.

The gene matrix generated from these reads initially included 6607 genes, but further filtering to remove rare transcripts resulted in 5620 genes for analysis. A total of 60 differentially expressed genes between the young and aged cohorts were identified at an FDR = 0.1, which we refer to as our “transcriptomic candidate list”. We found 42 of these genes were downregulated in the aged populations, while 18 were upregulated (Figure 3, Supplementary Figure 3, Supplementary Table 6).

**Figure 3:**
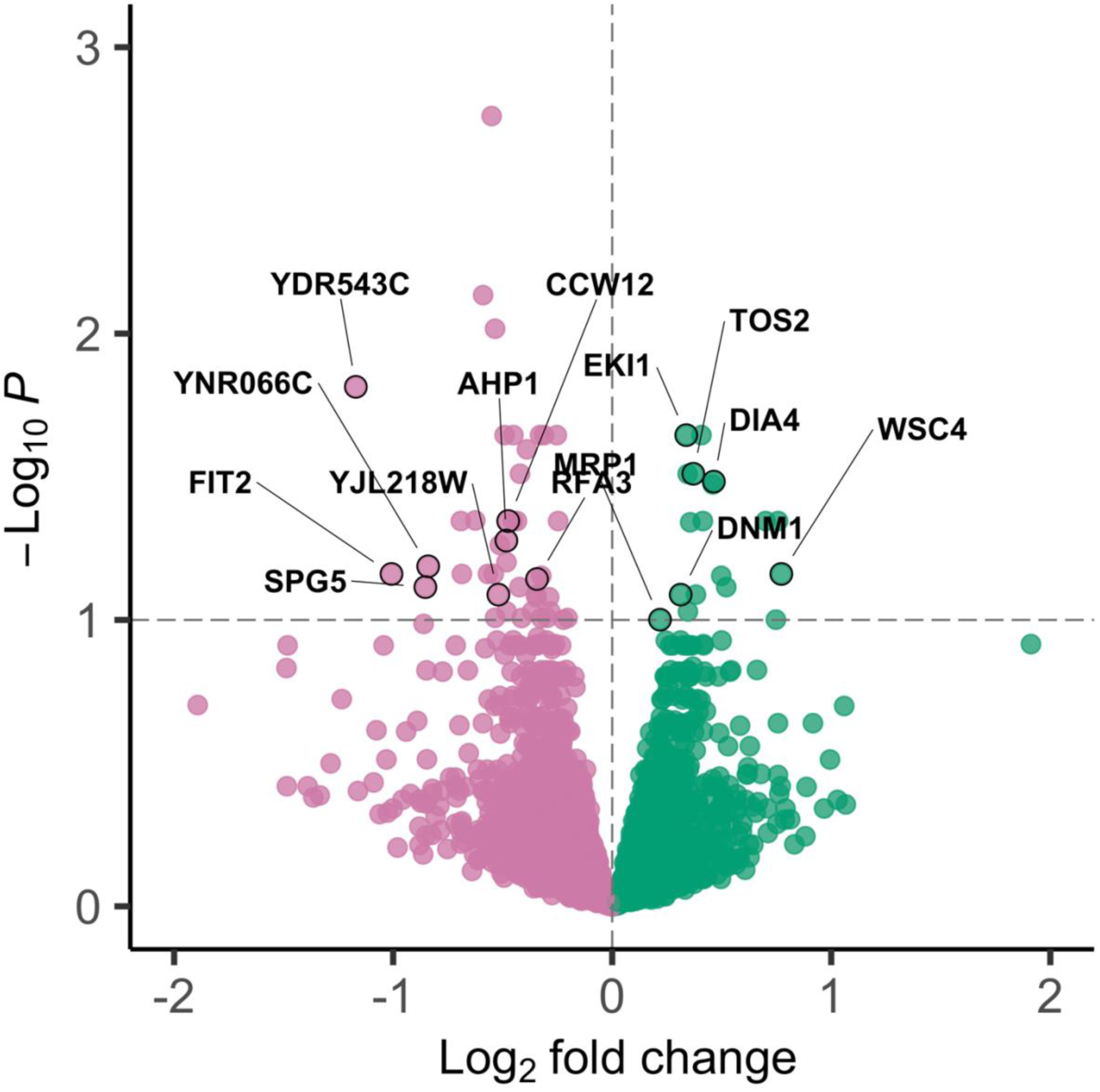
Volcano plot of differentially expressed genes. The horizontal dashed line represents significance threshold (adjusted p-value < 0.1). Green points represent genes that are upregulated in the aged cohorts (log_2_ fold change > 0), and pink points represent genes that are downregulated in the aged cohorts (log_2_ fold change < 0). Genes labelled in the plot (n=14) were identified as candidates in both the genomic and transcriptomic analyses (Table 1).

Within our transcriptomic candidate list, *YDR543C* had the greatest log_2_ fold change (approximately 125% reduction in expression in aged cohorts); however, *YDR543C* also had the lowest normalized expression count than any other feature at 20.9; by comparison, the mean expression value was 9154.16 and the median was 1180.323. The second highest change in expression was *FIT2*, with approximately 101% average decrease in expression in the aged cohorts compared to the young. *SPG5* and *YNR066C* also showed large reductions in expression, 81% and 79%, respectively. *WSC4* has the largest increase in expression in aged cohorts, at 41% (Figure 3, Supplementary Figure 3). *YRO2* had the next highest increase in expression in the aged cohorts, also nearing 41%, followed by *NDT80* and *SPS4* at 40% and 38% (Supplementary Figure 3).

No GO terms were found to be significantly overrepresented within our transcriptomic candidate gene list, so we examined the GO terms directly to characterize functional patterns. We found a few functions shared among sets of differentially expressed genes. Within the 18 upregulated genes, several appear to play roles in DNA replication and/or repair (*POL12, CLB6, CDC45),* transcription, (*GCR1, NDT80, OPI1*), translation (*DIA4, MRP1, SUI2*), and cell cycle regulation (*CLB6, NDT80, SPS4, TOS2*). DNA repair (*ACT1, MAG1, RFA3, UBC13*) and cell cycle regulation (*ACT1, CWP1, IRC18, RFA3*) were also implicated among downregulated genes. Other terms associated with downregulated genes included protein modification, targeting, folding, and breakdown (*EMA19, SSA1, UBC13, VID24, CPR6, RFA3, RPN6*), lipid metabolism (*CYB5, ERG25, IPT1, TMA7*), ion transport and homeostasis (*ATX1, FIT2, ATX2, VMA10*), and cell wall organization and biogenesis (*ACT1, CCW12, CWP1, IRC18*).

### Comparing genomic and transcriptomic analyses

First, we identified 114 significant genetic variants that were located within 10 kb of a differentially expressed gene; these local variants were distributed among 14 of the 60 differentially expressed genes from our transcriptomic candidate gene list. The other 46 genes were not associated with any significant local variants. Of the 14 genes with local genomic variants, 8 have variants within 5kb, and 4 within 1kb. Only 2 of these genes included variants within the gene transcript itself. Using SNPeff, we investigated the predicted functional implications of each of these variants (Table 1). While most of these variants were synonymous or intergenic, there were a substantial proportion of nonsynonymous variants (including two of the variants within the *RFA3* transcript).

**Table 1:**
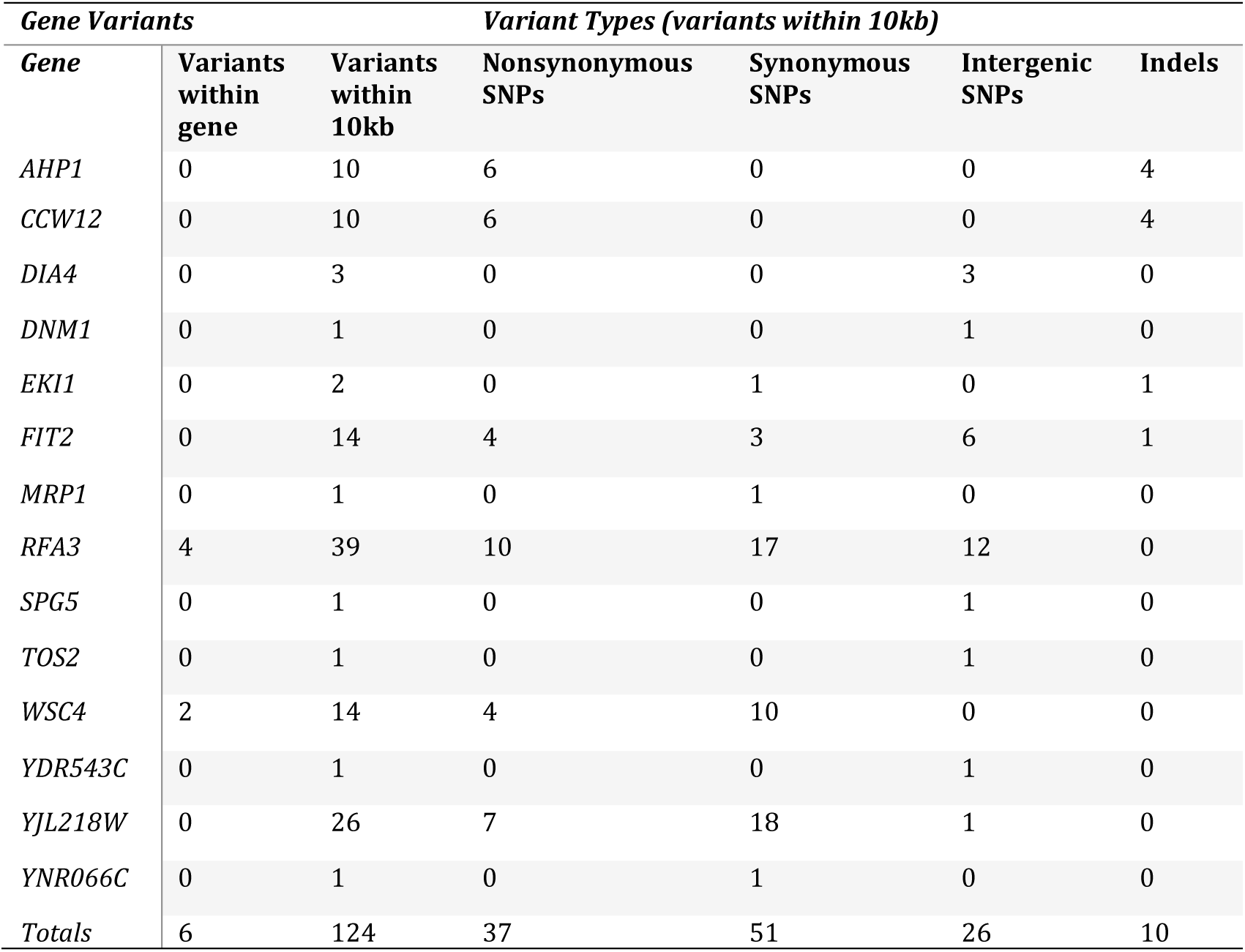
Differentially expressed genes with significant gene variants located within 10kb of the gene. Many upstream and downstream gene variants associated with these genes are also located within the coding regions of other genes, resulting in them being annotated as nonsynonymous and synonymous variants, rather than intergenic SNPs. However, six of these variants are located within the transcript of the differentially expressed gene: *RFA3* contains 2 nonsynonymous and 2 synonymous variants, while *WSC4* contains 2 synonymous variants.

The second method we used to search for potential *cis*-regulatory interactions was to investigate the representation of variant identities within the significant SNP data (e.g. nonsynonymous variants, synonymous variants, upstream gene variants, downstream gene variants) within our candidate genomic variant list and the annotated genome as a whole. These results should be interpreted with caution, since many of the annotations returned by SNPeff are loosely defined. Still, using a simulated Fisher’s exact test, we find that the distribution of annotation categories in our candidate list is statistically significant compared to annotations across the whole genome (p<0.001). Directly comparing the proportional representation of categories between our candidate gene list and the whole genome suggests that upstream gene variants and synonymous gene variants were underrepresented in our candidate gene list, while downstream gene variants and nonsynonymous variants were overrepresented compared to the whole genome (Figure 4).

**Figure 4:**
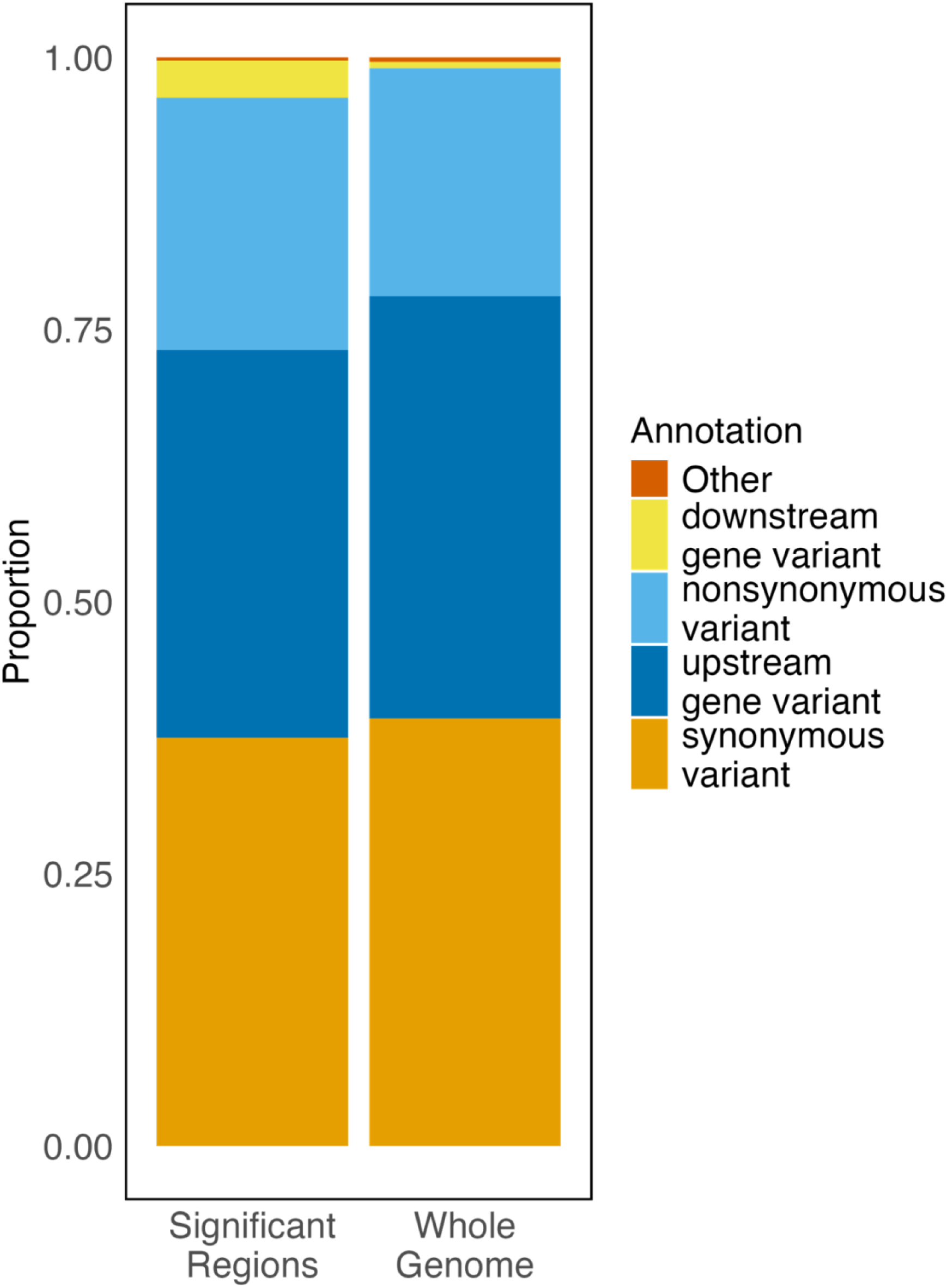
Annotation types for all significant SNP variants are shown for our significantly differentiated SNPs (left) compared to all known SNP variants annotated within SNPeff. If multiple annotations are present at a position, the first annotation was used.

Finally, we performed a GO-term analysis on the initial list of 14 genes that were candidates for local *cis-*regulatory interactions (p<0.1, FDR correction). We returned two terms, iron ion transport and iron coordination entity transport (both corresponding to the genes *FIT2* and *DNM1*).

## Discussion

We used bud scar staining and FACS to separate twelve paired replicates of aged and young populations for genomic and transcriptomic comparisons. To detect heritable age-associated variation, we sequenced sorted replicates after a short period of growth rather than immediately after sorting to standardize for growth stage. While our sorting approach achieved modest differences in replicative age phenotypes, genomic and transcriptomic data show clear differentiation between treatments. Haplotype analyses support that this differentiation is driven by selection rather than drift, as highly differentiated variants consistently map to specific founder haplotypes across independent replicates. Together, these findings demonstrate the presence of meaningful, heritable age-associated variation despite modest phenotypic divergence.

### Genomic differentiation implicates established and emergent aging pathways

Studies across diverse yeast strains have revealed consistent molecular hallmarks of cellular aging (reviewed by Janssens & Veenhoff 2016). Using existing genomic resources (SNPeff, SGD, UCSC Genome Browser), we found that many of our candidate genes fall within these well-established pathways, including protein homeostasis (e.g. *MNN* genes, *VID24*), nutrient sensing (e.g. *TOR2*), and DNA repair (e.g. *RFA3*). The recurrence of such canonical aging features validates our approach, while the appearance of less frequently cited but mechanistically linked genes highlights new avenues for exploring the polygenic landscape of aging.

Genes involved in protein synthesis, folding, targeting, and degradation were overrepresented in our dataset. A particularly striking pattern is the broad enrichment of glycosylation-related genes (e.g., *MNN* family genes, *PSA1*) alongside reduced expression of genes encoding abundant cell wall glycoproteins (*CCW12* and *CWP1*) in aged cohorts. Although cell wall staining could potentially bias detection toward cell wall-related processes, concordance between genomic and expression changes and prior literature suggest a biological basis. For example, CCW12 and CWP1 are both mannoproteins involved in new cell wall growth at bud sites (Ragni et al. 2011; Smits et al. 2006), and have shown binding affinity for WGA (Kung et al. 2009). If cells were upregulating WGA-binding proteins to escape the sorting protocol, we might expect both *CCW12* and *CWP1* show increased expression in our aged cohorts; instead, we find reduced expression in both genes. *CCW12* in particular makes up over 40% of cell wall glycoproteins and is essential for maintaining cell structure and morphogenesis (Ghanegolmohammadi et al. 2021), so its reduced expression implicates age-related shifts in cell wall composition that may impair structural integrity. While direct analogs are lacking in metazoans without cell walls, altered glycan composition and glycosylation patterns have also been associated with aging in mammals (Wu et al. 2025; Cindrić et al. 2021; Dall’Olio 2018; Krištić et al. 2014), supporting the evolutionary conservation of glycan metabolism as an aging modulator. The mechanistic links between the glycome and aging remain poorly understood (Palmer 2022; Miura & Endo 2016), but our results reinforce its relevance.

Consistent with prior work linking vacuolar dysfunction to declines in proteostasis and lifespan (Chen et al. 2020; Ghavidel et al. 2018; Hughes & Gottschling 2012), we also observed differentiation in genes involved in protein degradation (*VID24*, *VMA10*). *WSC4* (also called *YHC8*), one of only two genes implicated in both genomic and transcriptomic analyses, encodes a stress-response factor believed to influence protein targeting to the endoplasmic reticulum (Mamoun et al. 1999). These results support the hypothesis that maintenance of proteome integrity, via mechanisms mitigating the accumulation of misfolded or aggregated proteins, is a target of selection for increased replicative age (Saarikangas & Barral 2015; Aguilaniu et al. 2003).

Genes involved in DNA replication and repair also showed consistent differentiation. Upregulated genes in the aged cohorts included those essential for DNA replication initiation (*POL12*, *CLB6*, *CDC45*) and genotoxic stress response (*OPI1, CDC45, SPS4*), whereas downregulated genes included those involved in DNA repair (*MAG1, UBC13, RFA3*). *RFA3*, which encodes a subunit of Replication Protein A involved in both repair and telomere maintenance, is the second of the two genes implicated in both genomic and transcriptomic analyses. While previous work found *RFA3* upregulated in old cells (Lesur & Campbell 2004), we found it to be downregulated. This discrepancy likely reflects distinct dynamics in a recombinant genetic background where polygenic variation and interactions can emerge.

Metal ion homeostasis, particularly of iron, also emerged as a potential contributor to lifespan variation. Genes involved in iron and manganese transport and storage (*FIT2, ATX1, ATX2*) showed consistent expression differences. *FIT2*, located near highly differentiated genomic variants, has been linked to increased survival following DNA damage (Chen et al. 2020). These results suggest that metal ion availability influences DNA repair capacity in aging cells, potentially through interactions with polyamines such as spermine, which can stabilize DNA structures and modulate chromatin accessibility (Pedreño et al. 2005). This result highlights that natural genetic variation in metal ion regulators may therefore contribute to differences in replicative lifespan among yeast genotypes.

We also identified *TOR2*, a central component of the mTOR pathway and part of both TORC1 and TORC2 complexes (Kapahi et al. 2010), among our top genomic candidates. The TOR/mTOR pathway is one of several nutrient sensing pathways that has been linked to aging (Fontana et al. 2010), and has specifically been tied to replicative lifespan in yeast (Kaeberlein et al. 2005). Inhibition of TORC1 has been linked to lifespan extension (Seo et al 2024) and prevention of vacuolar fragmentation (Michaillat & Mayer 2013; Stauffer & Powers 2015, 2017), reinforcing its conserved role in aging. TOR (and DNA repair pathways) also play an important role in cell cycle regulation, which is inherently linked to replicative aging in yeast, as checkpoint mechanisms assess genomic integrity before division. Another candidate, *SWE1* (*Wee1*) regulates the G2/M transition and cell polarity, but has not been previously linked to aging. Our data suggest that *SWE1* may influence lifespan by modulating asymmetric inheritance of damaged cellular components, a process critical to rejuvenation in budding yeast (Lengefeld et al. 2017). Collectively, these results highlight diverse molecular pathways, including cell cycle regulation, proteostasis, glycan metabolism and metal ion homeostasis, as interacting components of the polygenic architecture of replicative aging.

Unlike expression-profiling studies of isogenic strains (e.g. Hendrickson et al. 2018), or temporal analyses of single lineages (Janssens & Veenhoff 2016), our use of recombinant populations allowed us to interrogate natural genetic variation underlying extended replicative lifespan. The overlap between our findings and known aging hallmarks validates our approach, while the discovery of novel contributors emphasizes the power of studies of standing genetic variation to deepen our understanding of complex traits.

### Regulatory mechanisms underlying polygenic variation for lifespan

Having characterized coding-region differentiation, we next asked whether noncoding or regulatory variation contributed to aging-associated differences. While most candidate variants (∼58%) occurred in genic regions, a substantial fraction were intergenic. Because RNA regulatory networks can influence aging (Montano & Long 2011; Čáp & Palková 2024), we searched for enrichment in regions involved in RNA surveillance but found none linked to canonical regulatory molecules (miRNAs, lncRNAs, snoRNAs). We did, however, detect age-associated upregulation of *AIR2,* a component of the TRAMP nuclear surveillance complex, consistent with regulation of RNA degradation mechanisms rather than noncoding RNA sequence variation. Notably, RNA exosome activity, including *AIR2*, has been implicated in maintaining cell wall integrity through regulation of protein glycosylation (Novačić et al. 2021).

Among the 191 unique genomic features implicated in our genomic and transcriptomic analyses, nine are annotated as “dubious ORFs”, which are unlikely to encode functional proteins. While these could represent artifacts or misannotated protein-coding genes, prior studies suggest some act as regulatory RNAs (Havilio et al. 2005). One of the ORFs we identified, *YDR543C*, lies within a telomeric region and may reflect altered transcription of telomeric RNAs (Aguado et al. 2020; Zeinoun et al. 2023). However, expression of this gene was very low and should be interpreted with caution. No other significant differences in expression were seen in telomeric regions. Overall, we find limited evidence that noncoding sequence variation directly drives aging-related regulatory differences between cohorts.

If local *cis*-regulatory interactions were a dominant mechanism promoting late life survival, we would expect substantial overlap between genes implicated in our genomic and transcriptomic analyses, or an enrichment of genetic variants in noncoding regions in close proximity to differentially expressed transcripts. To test this, we examined the distribution of significant genomic variants within 10, 5, and 1 kb of differentially expressed genes. Only 14 differentially expressed genes (24%) had nearby variants, with fewer under stricter thresholds, suggesting limited *cis*-regulatory influence.

Among significant candidate SNPs, upstream gene variants were underrepresented and downstream gene variants were overrepresented relative to genome-wide expectations. This pattern could indicate that enhancer-like or *trans*-regulatory sequences in downstream regions contribute more strongly to expression variation than promoter or more proximal variants (Savinkova 2009; Wittkopp & Kalay 2011). Given evidence that *trans*-regulatory interactions are more common in yeast (Siddiq & Wittkopp 2022), our data support the hypothesis that *trans*-acting and post-transcriptional mechanisms, rather than local sequence variation, underlie most regulatory divergence associated with replicative aging.

## Conclusions

While identifying genetic variants associated with aging is valuable, their biological relevance must be interpreted within regulatory contexts. Integrating genomic and transcriptomic data from the same populations enables inference of such relationships, revealing both shared candidates and broader regulatory trends that would be invisible using a single data type. Here, this approach clarified spatial relationships between differentially expressed genes and nearby variants, and provided a framework for linking genomic variation to transcriptional outcomes associated with aging.

Future research could extend this framework through iterative cycles of sorting and experimental evolution to select for progressively longer-lived populations, allowing tests of whether aging-related signals remain stable or shift over time. Incorporating additional -omic layers, such as epigenetic, proteomic, and metabolomic analyses, could further refine our understanding of how natural genetic variation shapes aging. Epigenetic modifications, for instance, may contribute to aging on short evolutionary timescales, but over how many generations or cycles of selection might those be sustained? Likewise, investigating proteomic and metabolic profiles could provide deeper insights into the functional consequences of age-associated genetic changes, particularly in relation to glycan metabolism and protein composition.

While our study was limited in its ability to isolate extremely old cells, we identified unambiguous signals from a comparison of populations following a single instance of bud-scar sorting into young and mid-aged cohorts. Additionally, haplotype data revealed that the allelic shifts observed in our aged replicates originated from three distinct ancestral haplotypes; this suggests that our approach of using genetically diverse populations of yeast allowed for the discovery of more complex adaptive responses that could not be identified from more traditional isogenic designs. Extending this approach across multiple age cohorts and selection cycles may ultimately disentangle how naturally-occurring variants and gene expression patterns jointly influence replicative lifespan.

## Methods

### Creation of experimental replicates

All experimental populations used in this study were derived from the “4S” yeast population described by Phillips et al. 2021. Briefly, this population was created by combining 4 haploid strains from the barcoded SGRP collection (YPS128, Y12, DBVPG6765, and DBVPG6044; Cubillos et al. 2009). Linder et al. 2020 modified these strains to enable easy crossing and diploid recovery (*MATa* strains have genotype *ycr043C::NatMX, ura3::KanMX-barcode, ho Δ,* and *MATα* strains have genotype *ycr043C::NatMX, ura3::KanMX-barcode, ho Δ*). After initial crossing, the 4S base population was subsequently subjected to 12 iterations of further outcrossing to maximize genetic diversity and allow for some domestication to laboratory conditions (Burke et al. 2020; Phillips et al. 2021).

The 4S population was revived from −80°C storage and 1 mL was plated as a lawn of cells on YPD media containing both 300 μg/mL hygromycin B (HYG) and 100 μg/mL nourseothricin sulfate (NTC), to ensure that only diploid cells (containing both mating types and both resistance cassettes) grew. The lawn of cells was broadly sampled with a wooden applicator and the resulting cellular material was used to inoculate 10 mL of YPD; this process was repeated to create 12 biological replicates. Each replicate was grown for 9 days at 30°C/200rpm to generate a distribution of cells with a range of replicative ages. This length of time was extended from the time period used for initial bud scar quantification (see Supplementary Methods) to allow more aged cells to accumulate in the population.

### Cell sorting and generation of paired age-cohorts

Prior to the experiment, we performed fluorescence imaging on sorted cells to confirm that the sorting process effectively separates aged cells from young cells (see Supplementary Methods, Supplementary Figures 4-5, Supplementary Table 7). After 9 days of aging, each replicate was standardized to a concentration of approximately 10^7^ cells/mL using an optical density measurement at 600nm (an OD_600_ of 1.0, +/- 0.1). Cell bud scars within each replicate were fluorescently stained with wheat-germ agglutinin Alexa fluor 488 (WGA CF488A; Biotium) per manufacturer instructions and sorted using a SONY SH800 cell sorter. A “main gate” was created to filter out particles of unusual size (e.g. large clumps of cells) that appeared as outliers based on a visual inspection of the scattered light signals during sorting setup; this gate typically retained ∼80-95% of particles. This gate was applied to both young and aged replicates. In addition to the main gate, the “aged” cohort for each replicate consisted of cells sorted from within the top 85-100% of fluorescence values while the “young” cohort consisted of cells sorted from within the bottom 50% of fluorescence values. Sorting of each replicate was stopped once it reached a count of 250,000 cells.

The 24 sorted fractions (12 pairs of young and aged replicate-cohorts) were transferred to 10 mL of YPD media and allowed to reach mid-log-phase growth before being archived to reduce potential noise from growth-phase-specific gene expression. All samples were collected for RNA sequencing when OD600 values ranged between 4 and 7, as growth-curve data suggest these values best predict mid-log phase. Approximately 1 million cells were collected from each replicate-cohort and immediately spun down. They were then preserved in 800 µL Trizol, which killed cells while preserving the RNA present in the sample to capture gene expression at the time of preservation. Each replicate preserved in Trizol was then homogenized by beadbeating for 30 sec prior to storage at −80°C. Samples for DNA sequencing were collected immediately following sample collection for RNA; 1 mL of culture was transferred directly to a cryotube and immediately frozen at −80°C.

### Genomic sequencing and analysis

Frozen samples were thawed and prepped using the Gentra Puregene Yeast/Bact Kit (Qiagen) following the manufacturer’s protocol. Genomic DNA libraries were prepared using the Illumina Nextera Library Prep Kit (Illumina), with some modifications to the protocol to increase throughput (Baym et al. 2015). All samples were dual-indexed and pooled for sequencing, with a size-selection step for 350-550 bp using a Qiagen MinElute Gel Extraction kit. The genomic DNA library was sequenced with the NextSeq 2000 (P2, PE150 reads) at the Center for Quantitative Life Sciences (CQLS) at Oregon State University.

We used a GATK-based variant calling pipeline (Phillips et al. 2021) to produce a merged VCF file containing all single-nucleotide polymorphisms (SNPs) and short insertion-deletion polymorphisms (indels) identified across all replicates and cohorts. Raw fastq files were aligned to the *S. cerevisiae* S288C reference genome (R64-2-1) using GATK v4.0 with standard best practices workflows and default filtering options (Van der Auwera & O’Connor 2020). The merged VCF file was converted into tables of variant frequencies by extracting the AD (allele depth) and DP (unfiltered depth) fields for all variants passing quality filters; the former field was used as the alternate allele count observed at a presumed variant, and the latter was used as the total coverage observed at that site. The VCF file was also used as an input for SnpEff v4.3 (Cingolani et al. 2012) to extract potential functional effects of individual SNPs. This variant frequency table was further filtered for minimum coverage values of 20 and maximum coverage of 1000 at every site in each replicate-cohort, and a minor allele frequency of at least 5% across the dataset. Differences in allele frequencies between the young and aged pairs of each replicate were evaluated via Cochran-Mantel-Haenszel (CMH) tests using the pool-seq package in R (Taus et al. 2017). The resulting *p*-values were log-transformed and a Bonferroni multiple testing correction (*α* = 0.05) was used to generate a genome-wide significance threshold. The SNPs and indels exceeding this threshold make up our “candidate genomic variant list”. This list was used to identify potentially enriched Genome Ontology (GO) terms through the Saccharomyces Genome Database Gene-Ontology Term Finder (Ashburner 2000; The Gene Ontology Consortium et al. 2023) with an FDR correction.

Using genome sequences from the four founder strains (Wing et al. 2020), we were able to estimate the most likely founder genotype at each position along the genome given our observed SNP frequencies. SNPs within a 5-kb sliding window (using a 2-kb step size) were used to estimate haplotype frequencies and the average divergence (“D”) between founder haplotypes across the genome as described in (Burke, King, et al. 2014; Burke, Liti, et al. 2014) The D statistic is the mean absolute difference in allele frequency between a haplotype’s frequency in the young cohort and its frequency in the aged cohort, averaged across all replicates. These values were plotted against expected (normally distributed) values to identify regions of unusually high divergence. We assessed signatures of selection (rather than genetic drift) by comparing the genome-wide SNP/indel and haplotype data, evaluating whether regions with elevated differentiation between young and aged replicates overlapped with regions of ancestral haplotype divergence consistently across replicates.

### Transcriptomic sequencing and differential gene expression analysis

Direct-zol RNA Miniprep Kits (Zymo Research) were used per manufacturer instructions to isolate total RNA. Each sample was treated with TURBO DNase enzyme (Invitrogen) and further purified and concentrated using the Clean and Concentrator kit (Zymo Research).

Samples were quantified with a Qubit fluorometer (3.0) and integrity assessed with an Agilent Bioanalyzer 2100 prior to library preparation using QuantSeq 3’ mRNA-Seq FWD kit (Lexogen, Vienna, Austria). The 24 barcoded libraries were sequenced as single-read 100-bp in an Illumina NextSeq2000 P1 flow cell at the CQLS.

We filtered the RNA-seq data using the recommendations provided by Lexogen QuantSeq for 3’ mRNA-seq. The bbduk tool was used to trim adapter sequences, polyA sequences, and bases with Phred scores below 20; we also removed reads shorter than 25 nucleotides. We used FastQC to check the quality of the filtered reads. After filtering, we performed genome alignment using the STAR alignment tool (Dobin et al. 2013), and used featureCounts (Liao et al. 2014) for quantification. The combined count matrix was filtered to remove genes with fewer than five mapped reads across all replicates and genes that have no expression (zero reads mapped) in 13 or more replicates. This retained 5620 genes for analysis.

We used Combat-seq (Zhang et al. 2020) to correct for a possible batch effect affecting pairs 1-6 and pairs 7-12, as these two groups were cultured and sorted on different days. We then performed differential expression analysis using DESeq2 (Love et al. 2014) in R. We compare expression (E) at each gene between aged and young cohorts for the *i^th^* pair (P) within the *j^th^* treatment (T), where the value of T_j_ is 0 for young and 1 for aged (Equation 4.1). *P*-values were corrected for false-discovery rate (FDR) and significance level assessed at FDR ≤ 0.1.

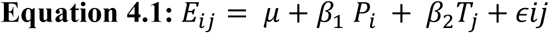

### Integrating genomic and transcriptomic analyses

Candidate gene lists generated from genome and transcriptome analyses were compared to determine if any genes were identified by both sequencing methods, as such overlap might implicate a *cis*-regulatory mechanism. We use three methods to identify potential *cis-*regulatory interactions using these two datasets: 1) we identified variants from our candidate genomic variant list located within 10kb of differentially expressed transcripts, 2) we analyzed predicted functional annotations of our genomic candidate variant list using SNPeff, and, 3) we performed a GO-term analysis on the genes that were implicated in both the genomic and transcriptomic candidate variant lists.

We first investigated gene variants that were near differentially expressed genes in our transcriptomic candidate variant list, and thus may be candidates for local *cis*-regulatory interactions. Local regulatory variation is commonly identified within promoter regions and the 3’ untranslated region (Ronald et al. 2005), and 10kb is often used as a threshold for detecting *cis*-regulatory interactions (e.g. Leach et al. 2007). Furthermore, SNPeff uses 5kb as a cutoff value for upstream and downstream gene variants (which should include most promoters and 3’ UTRs). However, many studies of *cis*-regulatory interactions set even stricter limits (e.g. 1 kb; (Shih & Fay 2021; Renganaath et al. 2020). Thus, we used 10 kb, 5kb, and 1kb as progressively conservative thresholds to detect evidence of potential local *cis*-regulatory interactions. This was done simply by mapping our differentially expressed genes to the genome using the Saccharomyces Genome Database and the UCSC Genome Browser, and finding each gene variant located within 10kb upstream or downstream of that gene in our candidate genomic variant list. This list also included variants located within differentially expressed genes.

Second, we looked at the predicted functional effects of significantly differentiated SNPs, comparing these to the predicted functional effects of all SNPs across the genome. We extracted the first annotation from SNPeff (sorted by deleteriousness) for each SNP that exceeded the significance threshold. We further grouped the SNP annotations so that any annotation category with a count of 1 or fewer in our candidate genetic variant list was grouped as “other” for both our candidate SNP list and the whole genome annotation data, leaving us with five functional annotation categories (downstream gene variants, upstream gene variants, nonsynonymous variants, synonymous variants, and “other”). A Fisher’s exact test was used to assess whether the proportion of SNPs assigned to each annotation category in our significant data differed from all annotated SNPs in the genome.

Finally, genes that were implicated in both the genomic and transcriptomic candidate gene lists were used for an additional GO-term enrichment analyses through the Saccharomyces Genome Database Gene-Ontology Term Finder (p<0.1 with an FDR correction).

## Supporting information

Supplementary Results

## Acknowledgements

We would like to thank Jeff Bishop and Adam Fries at the GC3F facility at the University of Oregon for their assistance with cell sorting and microscopy techniques, respectively.

## Data Availability Statement

Raw sequencing data have been deposited in the NCBI Sequence Read Archive under accession **PRJNA1368902,** *<u>and will be released upon publication</u>*. Code and processed data used for all analyses will be available in **a GitHub repository** (https://github.com/Katie-McHugh/regulation_of_aging/tree/Dryad). Additional datasets, including raw images and intermediate data files, will be archived with **Dryad** (DOI: 10.5061/dryad.34tmpg4x4) *<u>and made</u> <u>publicly available upon acceptance of the manuscript</u>*.

