## Supplementary Methods for "Age-specific genomic and transcriptomic variation reveals limited evidence for *cis-*regulatory interactions modulating aging in *Saccharomyces cerevisiae*"

Bud Scar Sorting and Quantification

*Sorting for age quantification*

The 4S population and each of its four isogenic founders (DBVPG6765, DBVPG6044, Y12, and YPS128) were revived from -80°C storage. The 4S population was revived as a diploid and plated on YPD containing both NTC and HYG. The four isogenic strains were revived as haploids, with *MATa* haploids grown on a YPD plate containing NTC and *MATα* haploids grown on a YPD plate containing HYG. The two mating types for each strain were replica plated on YPD plates containing both HYG and NTC to select for mated diploids. To generate five biological replicates for each population, a single colony of each isogenic strain was selected and used to inoculate 10 mL liquid YPD. For the 4S population, the lawn of cells on the plate was broadly sampled with a wooden applicator (to include as much of the genetic variation in the population as possible) and the resulting cellular material used to inoculate 10 mL of YPD. Sampling was done in groups of 2-3 replicates (with each group containing a mix of replicate populations) due to time limitations of cell sorting. These cultures were incubated in a shaking incubator at 30°C/200 rpm in 50mL conical vials. Each vial was incubated for 6.5 days to generate a wide distribution of replicative ages.

After aging, measurements of optical density at 600nm (OD600) were used to standardize each replicate to a concentration of approximately $10^7$ cells/mL (an $OD_{600}$ measurement of 1.0, +/- 0.1. Replicates were fluorescently stained with wheat-germ agglutinin Alexa fluor 488 (WGA CF488A; Biotium) per manufacturer instructions and sorted using a SONY SH800 cell sorter. A "main gate" was created to filter out particles of unusual size (e.g. large clumps of cells) that appeared as outliers based on a visual inspection of the scattered light signals during sorting setup; this gate typically retained ~80-95% of particles. In a preliminary test, the fraction of cells with the highest fluorescence values (4% +/- 2%) was found to be composed largely of auto-fluorescent cells, and subsequent dilution plating of these fractions revealed that most cells did not survive or produce colonies. Additionally, the average replicative age of surviving cells in this fraction was no higher than the average replicative age of a matched "aged" fraction that excluded the top 4% of cells (data not shown). Thus, this top fraction was excluded from the preliminary sort to reduce the proportion of dead cells in our aged cohorts. The lowest 78% of fluorescing cells (+/- 4%) were sorted into what we designated the "young" fraction, and cells with fluorescence ranging between these values were sorted into what we designated the "aged" fraction. A total of $10^6$ cells were sorted into each of the two fractions (aged and young) at a concentration of approximately 350,000-370,000 cells/mL. Approximately 400,000-500,000 cells (estimated by volume) from each sorted culture were spun down for 5 minutes at 14,000 rpm and resuspended in 20 uL of supernatant for imaging on an Olympus DeltaVision microscope at 600x magnification. Images were created using Z-stacking to generate multiple planes of view in each image so that bud scars would be visible regardless of their position on the cell. Images were uploaded to ImageJ for bud scar counting. For each age/replicate combination, approximately 24 images (+/-8, with one replicate of only 10 images) were saved for subsequent counting; the range is a result of images being excluded due to quality concerns. This typically resulted in 50-120 cells for counting in each replicate, with a minimum of 47 cells being counted. Of these, all replicates included at least 30 living cells except one (with 17 living cells). Unfortunately, the resolution achieved during fluorescence imaging made it difficult to distinguish bud scars in highly fluorescent cells with greater than 12-14 bud scars; however, the median replicative lifespan of all of the isogenic strains has been shown to range from 24-28 divisions (Kaya et al. 2021), and with a doubling time around 1-2 hours, cells easily could have achieved this many divisions during our experiment. Thus, the number of bud scars that could be confidently identified was recorded as a conservative estimate. Additionally, in some cases it was difficult to tell if a highly fluorescent cell was very old or dead; in this case, the cell was not counted.

After the number of bud scars on each cell in each image had been recorded, Shapiro tests were used to assess the normality of the distribution of cell ages. Parametric and non-parametric pairwise comparisons were used to assess differences in mean bud scar count between the young and aged sorted fractions within each population as appropriate.
