## Supplementary Results for "Age-specific genomic and transcriptomic variation reveals limited evidence for *cis-*regulatory interactions modulating aging in *Saccharomyces cerevisiae*"

Low fluorescence sorted fractions (young cohorts) had fewer bud scars (and reduced replicative age) on average compared to their paired high fluorescence (aged) counterparts (see Supplementary Figure 5). This pattern was consistently observed across each of the five tested populations, with only one population not achieving statistical significance (p-values for each population shown in  Supplementary Table 7).
