## Supplementary Table Captions for "Age-specific genomic and transcriptomic variation reveals limited evidence for *cis-*regulatory interactions modulating aging in *Saccharomyces cerevisiae*"

**Supplementary Table 1: Genome-wide coverage values for each replicate.** Data is shown for both the SNP and indel datasets.

**Supplementary Table 2: Candidate genomic variant list.** Genome variants that are significantly differentiated between young and old replicates. Table includes both SNPs and indels from the nuclear and mitochondrial genomes ($p<0.05$ with Bonferroni correction).

**Supplementary Table 3: GO-terms returned for significant results.** These GO-terms were overrepresented among our candidate variant list for the genome or within genes that were present in both genomic and transcriptomic variant lists (listed in Table 1). GO-term analysis was conducted using the Saccharomyces Genome Database Gene-Ontology term finder with an FDR correction and $p<0.05$. No GO-terms were returned for the list of significantly differentially expressed transcripts ($p<0.1$, FDR correction).

**Supplementary Table 4: Most significant peaks of haplotype differentiation.** Elevated and reduced haplotypes for each peak are shown, along with genes that harbor genetic variants within these regions.

**Supplementary Table 5: Read and base counts for each replicate throughout each step of our transcriptomic analysis.** Read and base counts are shown after initial sequencing, filtering (bbduk), alignment (STAR), and quantification (featureCounts).

**Supplementary Table 6: All differentially expressed genes ($p<0.1$, FDR correction) in the nuclear genome.** This table shows the location of each transcript within the genome, results data generated from the DEG analysis, and the relationship of each transcript to the closest gene variants identified in the genome-wide analysis.

**Supplementary Table 7: Bud scars in young and aged sorted fractions.** The number of bud scars counted per cell in each population was averaged across biological replicates. P-values in the table come from within-population comparisons as evaluated by pairwise tests with t-tests (no shading) and Wilcoxon signed rank tests (shaded in gray), used as appropriate. Significant differences between young and aged cohorts within each population are indicated by an asterisk (*).
