## Supplementary Table 3 for "Age-specific genomic and transcriptomic variation reveals limited evidence for *cis-*regulatory interactions modulating aging in *Saccharomyces cerevisiae*"

| *Gene List* | *Category* | *Term* | *P-value* | *Associated features* |
|---|---|---|---|---|
| **Genome** | Component | cell periphery | 5.49E-07 | AIM44, HKR1, YHL026C, SIM1, FKS1, YPS6, CRH1, JEN1, HSC82, TOR2, HXT3, WSC3, FRE3, HIP1, THI7, BBC1, DAN4, TDH3, STE6, GTS1, AGA1, PHO90, TIR1, OPT2, YNL190W, SAC1, FIG2, TIR2, MTL1, SEC3, PGA3, ADH1, YBR067C, SUC2, NUM1, FLO11 |
| | | cell wall | 1.73E-03 | YNL190W, DAN4, FIG2, TDH3, AGA1, TIR1, YPS6, CRH1, SIM1, YBR067C |
| | | fungal-type cell wall | 1.73E-03 | AGA1, TIR1, CRH1, YPS6, SIM1, YBR067C, YNL190W, DAN4, FIG2, TDH3 |
| | | external encapsulating structure | 1.73E-03 | SIM1, CRH1, YPS6, YBR067C, AGA1, TIR1, FIG2, TDH3, YNL190W, DAN4 |
| | | plasma membrane | 1.10E-02 | MTL1, PGA3, THI7, ADH1, FRE3, HIP1, STE6, FLO11, TDH3, HKR1, AIM44, PHO90, JEN1, FKS1, OPT2, WSC3, HXT3, HSC82, TOR2 |
| | | golgi apparatus | 4.00E-02 | YKT6, MNN2, MNN5, MNN1, MNN4, STE6, SAC1, MNN11, YND1, GEA2, OPT2, SPF1 |
| | | golgi cisterna | 8.79E-02 | MNN11, SAC1, GEA2, OPT2 |
| | Process | protein glycosylation | 1.10E-02 | MNN2, MNN4, YND1, MNN1, MNN11, MNN5, ALG7, PMI40 |
| | | macromolecule glycosylation | 1.10E-02 | ALG7, PMI40, MNN11, MNN1, MNN5, YND1, MNN2, MNN4 |
| | | glycosylation | 1.30E-02 | ALG7, PMI40, MNN1, MNN11, MNN5, YND1, MNN4, MNN2 |
| | | glycoprotein biosynthetic process | 1.80E-02 | MNN5, MNN1, MNN11, PMI40, ALG7, MNN2, MNN4, YND1 |
| | | glycoprotein metabolic process | 4.30E-02 | YND1, MNN4, MNN2, PMI40, ALG7, MNN5, MNN1, MNN11 |
| | Function | NA | NA | NA |
| **Transcriptome** | NA | No Significant GO-terms | NA | NA |
| **Shared Genes** | Process | Iron Coordination entity transport | 3.13E-02 | DNM1, FIT2 |
| | | iron ion transport | 9.98E-02 | FIT2, DNM1 |
| | Component | NA | NA | NA |
| | Function | NA | NA | NA |

**Supplementary Table 3: GO-terms returned for significant results.** These GO-terms were overrepresented among our candidate variant lists for the genome and within genes that were present in both genomic and transcriptomic variant lists (“Shared Genes”, Table 1). GO-term analysis was conducted using the Saccharomyces Genome Database Gene-Ontology term finder with an FDR correction and $p<0.05$. No GO-terms were returned for the list of significantly differentially expressed transcripts ($p<0.1$, FDR correction).
