## Supplementary Table 4 for "Age-specific genomic and transcriptomic variation reveals limited evidence for *cis-*regulatory interactions modulating aging in *Saccharomyces cerevisiae*"

| *Peak* | *Chromosome* | *Elevated haplotype* | *Reduced haplotype* | *Associated features* |
|---|---|---|---|---|
| *A* | 5 | DBVPG6765 | DBVPG6044 | MNN1, MIT1,YND1, PMP2, WBP1, NOP16, IRC22, PMI40, TMA20, NUG1, TMA20, SEC3, SNR14, TIR1, YER010C |
| *B* | 10 | YPS128 | DBVPG6044 | MNN5, SWI3, SWE1, RFA3, RPS22A, ATG36, YJL181W, YJL182C, ELO1, ATG27, ATP12, YJL163C, YJL202C, ACO2, LAA1, RPS14B, YJL171C, SOP4, YJL182C, PHO90, UBP12, MNN11, YJL197C-A, RPS22A, CPS1 |
| *C* | 11 | Y12 | DBVPG6765 | PTK1, MNN4, TOR2, YKL202W, UBA1, TRP3, STE6, PEX1, SDS22, EAP1, EMC3, JEN1, CBT1, YKT6, URA1, DOA1, SAC1 |

**Supplementary Table 4: Most significant peaks of haplotype differentiation.** Elevated and reduced ancestral haplotypes for each peak are shown, along with genes that harbor genetic variants within peak regions.
