## Supplementary Table 7 for "Age-specific genomic and transcriptomic variation reveals limited evidence for *cis-*regulatory interactions modulating aging in *Saccharomyces cerevisiae*"

| ***Population*** | ***Origin*** | ***n=*** | ***p-value*** | ***Mean bud scars in young fractions*** | ***Mean bud scars in aged fractions*** |
|---|---|---|---|---|---|
| *4S* | Recombinant population | 5 | 0.00389* | 2.144 | 2.826 |
| *YPS128* | North American oak forest | 5 | 0.00091* | 2.006 | 2.756 |
| *Y12* | Japanese sake | 5 | 0.0625 | 2.115 | 3.044 |
| *DBVPG6044* | African palm wine | 4 | 0.00453* | 1.8 | 3.172 |
| *DBVPG6765* | European wine | 4 | 0.0284* | 2.143 | 3.235 |

**Supplementary Table 7: Bud scars in young and aged sorted fractions.** The number of bud scars counted per cell in each population was averaged across biological replicates. P-values in the table come from within-population comparisons as evaluated by pairwise tests with t-tests (no shading) and Wilcoxon signed rank tests (shaded in gray), used as appropriate. Significant differences between young and aged cohorts within each population are indicated by an asterisk (*).
