## Supplementary Figure 1 for "Age-specific genomic and transcriptomic variation reveals limited evidence for *cis-*regulatory interactions modulating aging in *Saccharomyces cerevisiae*"

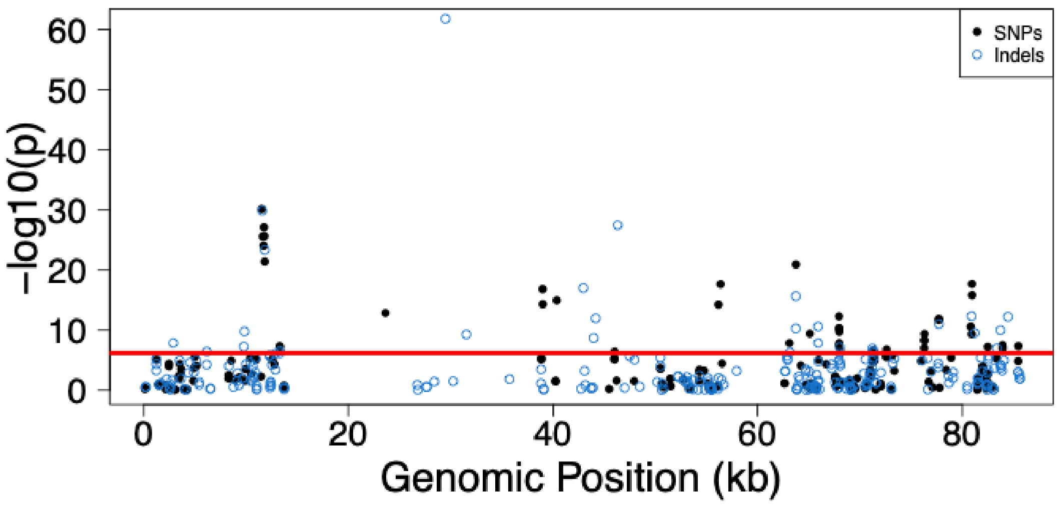


**Supplementary Figure 1:** Regions of genome differentiation within the mitochondria. SNPs are shown in black and indels are shown in blue. The red line shows a Bonferroni corrected α=0.05 threshold, generated using SNPs and indels from the whole genome.
