## Supplementary Figure 2 for "Age-specific genomic and transcriptomic variation reveals limited evidence for *cis-*regulatory interactions modulating aging in *Saccharomyces cerevisiae*"

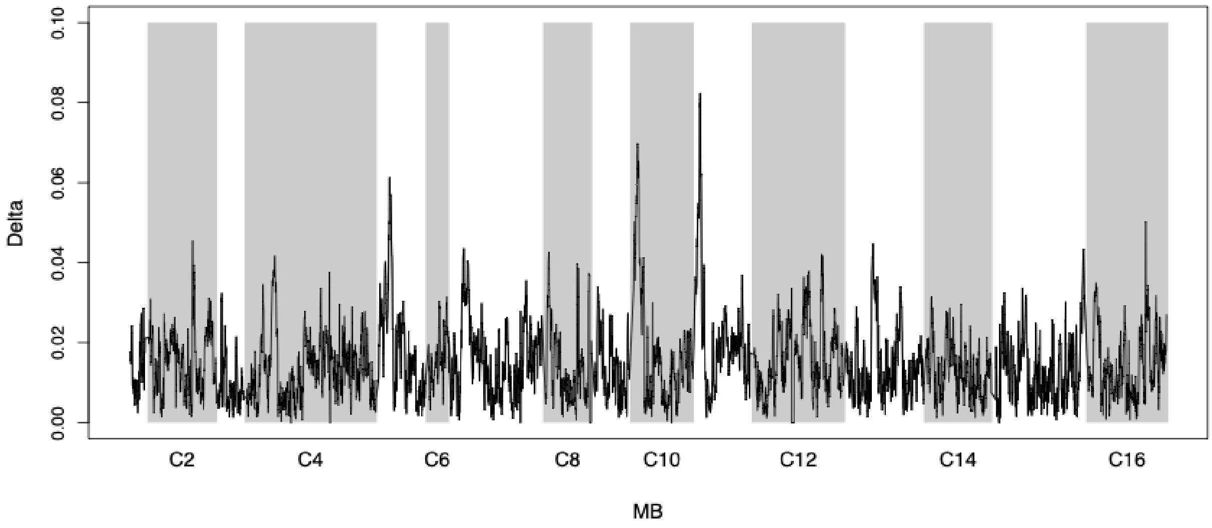


**Supplementary Figure 2:** Haplotype divergence ("D") across the genome. The average divergence between all ancestral haplotypes at each point in the nuclear genome is shown
