## Supplementary Figure 3 for "Age-specific genomic and transcriptomic variation reveals limited evidence for *cis-*regulatory interactions modulating aging in *Saccharomyces cerevisiae*"

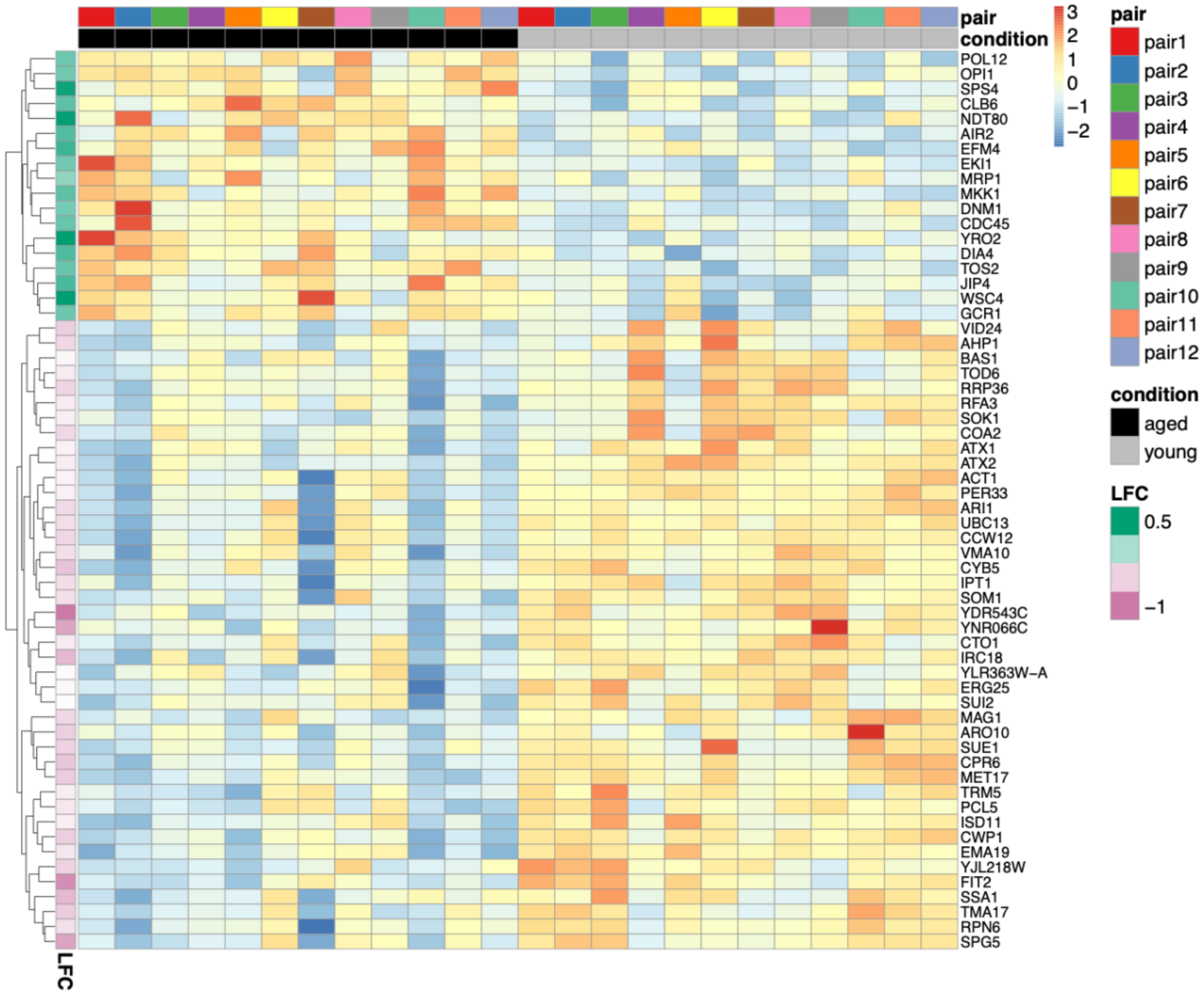


**Supplementary Figure 3:** Heatmap showing differential expression of significant genes ($p<0.1$). The top bar indicates the pairing structure of the data, and the second bar indicates the age of the replicate. The bar on the far left shows the average $\log_2$ fold change (LFC) of expression across replicates. Positive values of LFC (in green) indicate an increase in expression in the aged replicates.
