## Supplementary Figure 4 for "Age-specific genomic and transcriptomic variation reveals limited evidence for *cis-*regulatory interactions modulating aging in *Saccharomyces cerevisiae*"

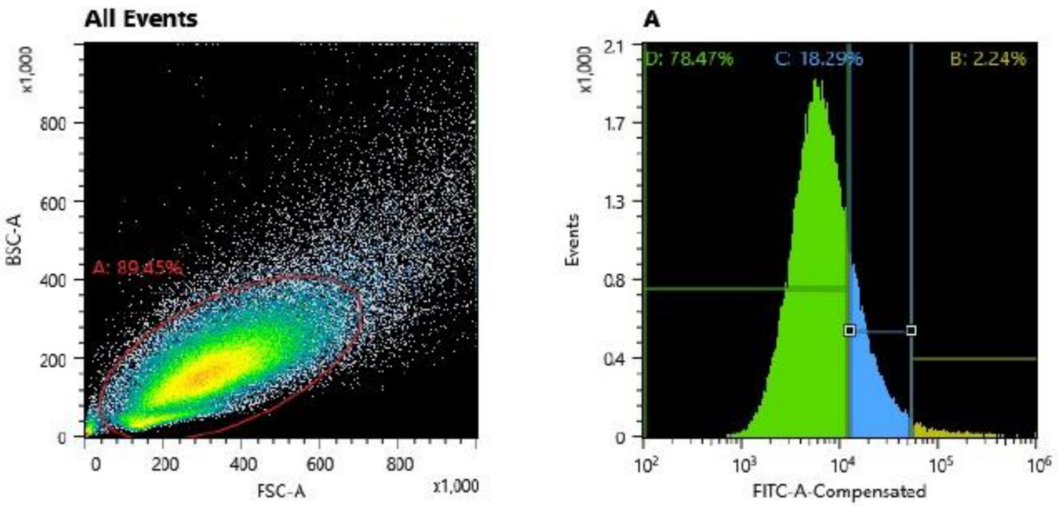


**Supplementary Figure 4: Example of the readout used to sort cells using FACS.** The figure on the left is a plot of the frontscatter (FSC-A) and backscatter (BSC-A). The red circle (A) indicates the main gate. The plot on the left shows only particles within the main gate, and depicts the number of particles (events) displaying different fluorescence intensities. Gate D (in green) includes lower-fluorescence cells sorted into our young cohort. Gate C (blue) contains higher fluorescence cells sorted into our old cohort. Gate B (yellow) was excluded during preliminary sorts due to the high concentration of dead cells; however, these cells were retained during the experimental sorts to avoid excluding very old cells.
