## Supplementary Figure 5 for "Age-specific genomic and transcriptomic variation reveals limited evidence for *cis-*regulatory interactions modulating aging in *Saccharomyces cerevisiae*"

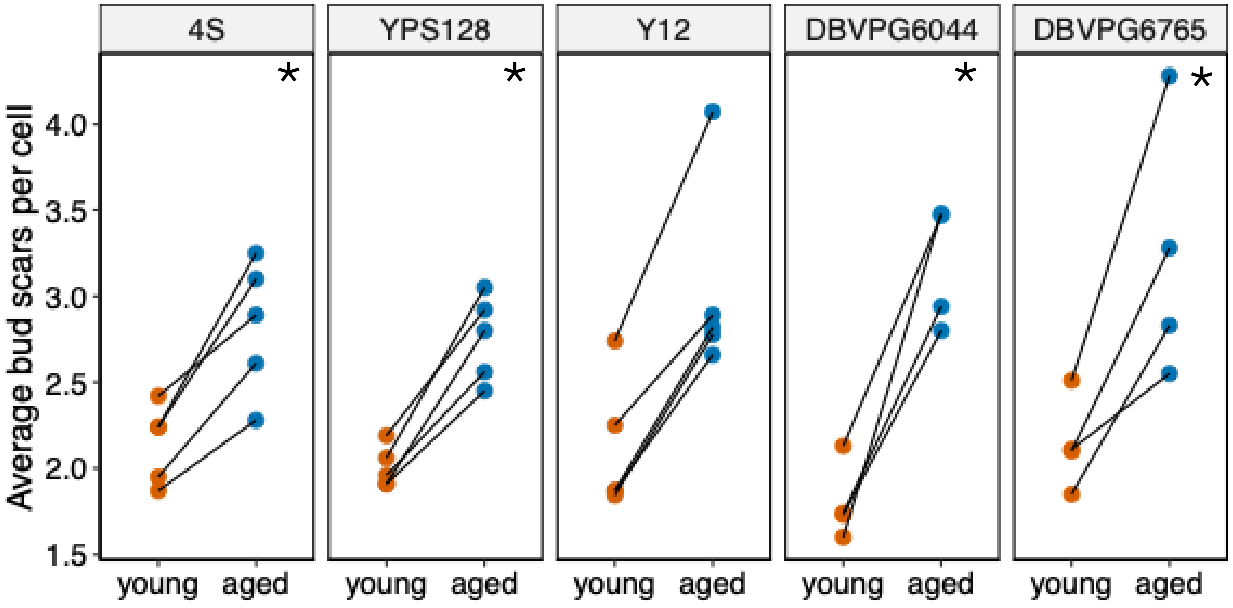


**Supplementary Figure 5: Bud scars in young and aged sorted fractions.** Red points (left side of each panel) indicate young and blue points (right) indicate aged sorted fractions, with lines between points indicating the paired nature of the data points. For the 4S, Y12, and YPS128 populations n=5 pairs of replicates. For the DBVPG6044 and DBVPG6765 strains n=4 pairs of replicates. Asterisks (*) indicate statistically significant differences.
